# Piezo1 and BK_Ca_ channels in human atrial fibroblasts: interplay and remodelling in atrial fibrillation

**DOI:** 10.1101/2021.01.21.427388

**Authors:** Dorothee Jakob, Alexander Klesen, Benoit Allegrini, Elisa Darkow, Diana Aria, Ramona Emig, Ana Simon Chica, Eva A. Rog-Zielinska, Tim Guth, Friedhelm Beyersdorf, Fabian A. Kari, Susanne Proksch, Stéphane N. Hatem, Matthias Karck, Stephan R Künzel, Hélène Guizouarn, Constanze Schmidt, Peter Kohl, Ursula Ravens, Rémi Peyronnet

**Affiliations:** Institute for Experimental Cardiovascular Medicine, University Heart Center Freiburg · Bad Krozingen, Medical Center - University of Freiburg, Germany; Faculty of Medicine, University of Freiburg, Germany; CNRS University Cote d’Azur laboratory Institut Biology Valrose, Nice, France; Spemann Graduate School of Biology and Medicine (SGBM), University of Freiburg, Freiburg, Germany; Faculty of Biology, University of Freiburg, Freiburg, Germany; G.E.R.N. Tissue Replacement, Regeneration & Neogenesis, Department of Operative Dentistry and Periodontology, Medical Center - University of Freiburg, Germany; CIBSS Centre for Integrative Biological Signalling Studies, University of Freiburg, Germany; Department of Cardiovascular Surgery, University Heart Center Freiburg · Bad Krozingen, Medical Center - University of Freiburg, Germany; Sorbonne University; Assistance Publique-Hôpitaux de Paris, GH Pitié-Salpêtrière Hospital, INSERM UMR_S1166; Cardiology department, Institute of Cardiometabolism and Nutrition-ICAN, Paris; Department of Cardiac Surgery, University of Heidelberg, Germany; Institute of Pharmacology and Toxicology, Faculty of Medicine Carl Gustav Carus, Technische Universität Dresden, Germany; Department of Cardiology, University of Heidelberg, Germany; and DZHK (German Center for Cardiovascular Research) partner site Heidelberg / Mannheim, University of Heidelberg, Germany

**Keywords:** Stretch-activated ion channels, mechano-sensing, heart, arrhythmia, non-myocytes, calcium, Slo1

## Abstract

**Aims:** Atrial Fibrillation (AF) is an arrhythmia of increasing prevalence in the aging population of developed countries. One of the important indicators of AF is sustained atrial dilatation, highlighting the importance of mechanical overload in the pathophysiology of AF. The mechanisms by which atrial cells, including fibroblasts, sense and react to changing mechanical forces, are not fully elucidated. Here, we characterise stretch-activated ion channels (SAC) in human atrial fibroblasts and changes in SAC-presence and -activity associated with AF.

**Methods and Results:** Using primary cultures of human atrial fibroblasts, isolated from patients in sinus rhythm or sustained AF, we combine electrophysiological, molecular and pharmacological tools to identify SAC. Two electrophysiological SAC-signatures were detected, indicative of cation-nonselective and potassium-selective channels. Using siRNA-mediated knockdown, we identified the nonselective SAC as Piezo1. Biophysical properties of the potassium-selective channel, its sensitivity to calcium, paxilline and iberiotoxin (blockers), and NS11021 (activator), indicated presence of calcium-dependent ‘big potassium channels’, BK_Ca_. In cells from AF patients, Piezo1 activity and mRNA expression levels were higher than in cells from sinus rhythm patients, while BK_Ca_ activity (but not expression) was downregulated. Both Piezo1-knockdown and removal of extracellular calcium from the patch pipette resulted in a significant reduction of BK_Ca_ current during stretch. No co-immunoprecipitation of Piezo1 and BK_Ca_ was detected.

**Conclusions:** Human atrial fibroblasts contain at least two types of ion channels that are activated during stretch: Piezo1 and BK_Ca_. While Piezo1 is directly stretch-activated, the increase in BK_Ca_ activity during mechanical stimulation appears to be mainly secondary to calcium influx *via* SAC such as Piezo1. During sustained AF, Piezo1 is increased, while BK_Ca_ activity is reduced, highlighting differential regulation of both channels. Our data support the presence and interplay of Piezo1 and BK_Ca_ in human atrial fibroblasts in the absence of physical interactions between the two channel proteins.

## 1. Introduction

Atrial Fibrillation (AF) is a supraventricular arrhythmia with increasing prevalence in countries with an aging population. Although AF is one of the most common cardiovascular causes of hospitalization,^1–3^ its pathophysiology is not fully elucidated, and it represents an unmet need for effective prevention and treatment. One hallmark of AF is its progressive nature. As AF becomes increasingly resistant over time to pharmacological or electrical attempts at conversion back to sinus rhythm (SR),^2^ atrial tissue undergoes pronounced remodelling.^2, 4^ Structural and functional changes involve cell electrophysiological and tissue morphological alterations. Whilst electrical remodelling of atrial cardiomyocytes is characterized by a shortening in action potential duration and of effective refractory period, as well as by impaired adaptation of these parameters to changes in heart rate,^5^ fibrosis – a prominent feature of AF-related structural remodelling – may in parallel contribute to slowing of conduction. The combination of short effective refractory period and slow conduction favours maintenance of AF *via* re-entry mechanism.^6^

Many of the risk factors for AF, *e.g.* heart failure, hypertension, or valvulopathies, are accompanied by mechanical overload of the atria.^7^ Since stretch enhances the susceptibility to AF induction,^8, 9^ it has been suggested, that mechanical overload may contribute to initiation and perpetuation of AF *in vivo*.^10–13^ In addition, acute stretch of control atrial tissue induces complex and regionally varying changes in action potential shape,^10^ and diastolic depolarization which can trigger extrasystoles.^9, 14, 15^ This ‘mechano-electric feedback’^16, 17^ requires cells to be able to sense their mechanical environment, and to translate this into an electrophysiologically relevant signal.

Ample evidence points to an essential role of stretch-activated ion channels (SAC) as mechano-sensors in cardiomyocytes (for reviews see ^18, 19^). SAC are also present and functional in human atrial fibroblasts,^20, 21^ but it is currently not known whether SAC function is altered in AF in human heart cells, especially in fibroblasts, which are key players in fibrosis. Therefore, the aim of this study was to compare SAC function in atrial fibroblasts from patients in SR and sustained AF. The cation-nonselective SAC Piezo1^22–24^ forms a plausible candidate, in line with recently reported Piezo1 effects on remodelling of non-cardiac tissues.^25^ A second candidate, the potassium-selective Ca^2+^-activated channel of large conductance (BK_Ca_), has been reported to respond to stretch, local Ca^2+^ concentration changes, TGF-β, and angiotensin II in several cardiac cell types.^26–30^ BK_Ca_ is further known to modulate fibroblast proliferation,^31^ a critical event during pathological tissue remodelling in AF. Both Piezo1 and BK_Ca_ have previously been detected in human atrial fibroblasts.^31–33^

In this study we report AF-related changes in Piezo1 and BK_Ca_ channel activity in human atrial fibroblasts, and establish functional interactions between the two channel types.

## 2. Material and Methods

### 2.1 Tissue collection

Tissue samples were obtained from the right atrial appendage of patients undergoing open-heart surgery at the University Heart Center Freiburg · Bad Krozingen. Patients were either in SR, or in sustained AF (which includes patients with persistent, long-standing persistent and permanent AF, defined according to ESC Guidelines).^34^ Tissue samples were processed by the Cardiovascular Biobank of the University Heart Center Freiburg · Bad Krozingen (approved by the ethics committee of Freiburg University, No 393/16; 214/18) or the Clinical Center of the Medical Faculty Heidelberg (approved by the ethics committee of the University Heidelberg, S-017/2013). Upon excision in the operating theatre, tissue was placed in room-temperature cardioplegic solution (containing in [mmol/L]: NaCl 120, KCl 25, HEPES 10, glucose 10, MgCl_2_ 1; pH 7.4, 300 mOsm/L) and immediately transported to the laboratory. Tissue was processed within 30 min of excision. Samples from 36 SR patients and 17 AF patients (mean age 64.8 ± 1.4 years [mean ± standard error of the mean, SEM], age range 38 - 83 years, 40 males, 13 females were used (Table 1). No significant age differences between the two groups were observed. Left atrial diameter of AF patients was larger, compared to SR patients. All patients gave informed consent prior to inclusion in the study, and investigations conformed to the principles outlined in the Declaration of Helsinki.

**Table 1:**
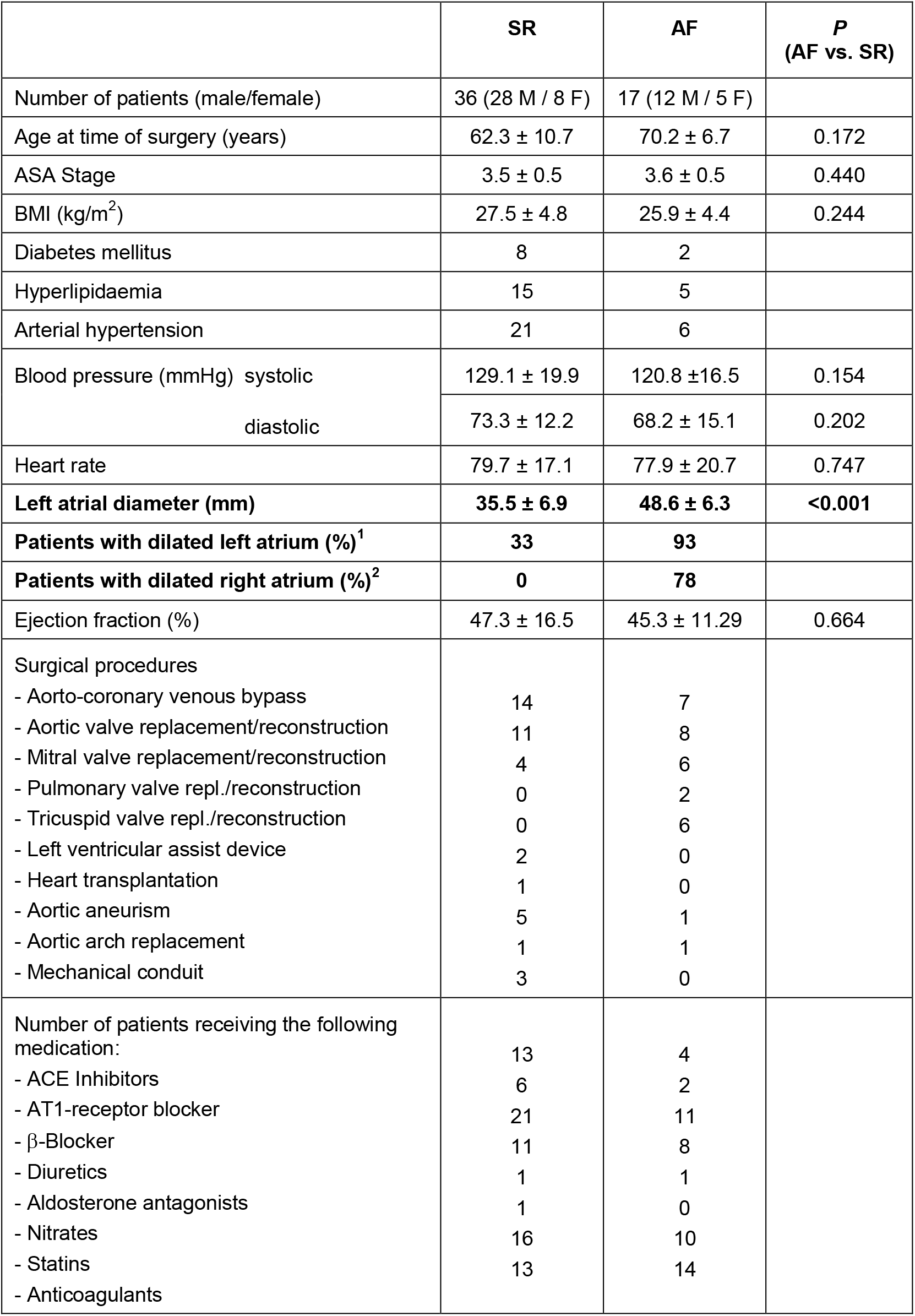
Patient characteristics. ASA: American Society of Anaesthesiologists, BMI: body mass index, ACE: Angiotensin 144 converting enzyme, AT1: Angiotensin II receptor type I; mean ± SD. ^1^information available for 145 24 SR and 14 AF patients; ^2^information available for 23 SR and 14 AF patients.

### 2.2 Cell culture

The size of the tissue samples was variable (50 to 200 mg). The epicardium and adipose tissue were carefully removed to avoid contamination with excess epicardial cells or adipocytes. The remaining myocardium was cut into blocks of about 1-4 mm^3^. Tissue chunks were transferred into a 6-well plate, each well containing 2 mL of Dulbecco’s Modified Eagle Medium (DMEM, Gibco, Germany), 10% foetal calf serum, and 1% penicillin/streptomycin (all Sigma-Aldrich, Germany), for incubation at 37°C in an atmosphere of air supplemented with CO_2_ to maintain 5% CO_2_. Culture medium was changed twice a week. Prior to use, the surface of culture plates had been abraded using a scalpel blade to favour tissue attachment and cell propagation. This so called “outgrowth technique”^35^ was used for functional experiments as it yields more reproducibly large numbers of viable cells, compared to enzymatic digestion.

After 7-10 days, cells started to migrate from the tissue chunks and reached ~80% confluency after 20-28 days, when they had to be passaged to preserve viability. For passaging, culture medium was removed, and cells were washed with pre-warmed 1% phosphate-buffered saline (PBS) solution, detached by adding 1 mL of 1% trypsin per 35-mm dish for 5-10 min. After addition of 2 mL of culture medium per dish, the suspension was transferred into a 15 mL Falcon tube and centrifuged at 333 × g for 5 min. The supernatant was carefully removed and discarded. The cell pellet was resuspended in 1 mL of pre-warmed culture medium; 10 μL of this cell suspension were counted in a Neubauer chamber. Then cells were seeded into culture flasks or dishes for further experiments. To achieve a uniform cell density, 25,000 cells were seeded per 35-mm dish. Cells were used for experimentation until passage 4 (*i.e*. for up to 6 weeks). For co-immunoprecipitation experiments, the recently developed^36^ human atrial fibroblast cell line HAF-SRK01 (HAF) was also used. Culture conditions for HAF and primary cultures were identical.

For reference purposes we also used atrial fibroblasts that were freshly isolated by enzymatic dissociation.^35^ In brief, the right atrial tissue samples were placed into Ca^2+^-free modified ‘Kraftbrühe’ solution (in mmol/L, KCl 20, K_2_HPO_4_ 10, glucose 25, D-mannitol 40, K-glutamate 70, ß-hydroxybutyrate 10, taurine 20, EGTA 10, pH 7.2) supplemented with albumin (0.1%).^37^ Fat and epicardial tissue were removed, and the remaining tissue was cut into pieces of 1-4 mm^3^, followed by rinsing for 5 min with Ca^2+^-free solution supplemented with taurine. The solutions were oxygenated with 100% O_2_ at 37°C and stirred. For digestion, tissue aliquots were transferred for 10 min into a Ca^2+^-free solution (in mmol/L, NaCl 137, KH_2_PO_4_ 5, MgSO_4_(7H_2_O) 1, glucose 10, HEPES 5, pH 7.4) supplemented with taurine, albumin (0.1%), collagenase type V (200 U/mL) and proteinase XXIV (5.4 U/mL). The Ca^2+^ concentration was then increased to 0.2 mmol/L and the tissue was stirred for additional 20-30 min. An additional 10 min of incubation in the presence of collagenase was then performed to release the first cells. This step was repeated until complete digestion of the tissue was accomplished. Collected suspensions were centrifuged (7 × g for 2 min) to separate cardiomyocytes (in the pellet) from non-myocytes (in the supernatant). Cardiomyocytes were resuspended in 250 μL of lysis buffer (RLT, Qiagen, Germany) mixed with 10 μL of β-mercaptoethanol and frozen at −80°C. Non-myocytes where centrifuged at 260 × g for 5 min. Cell pellets were resuspended in lysis buffer and frozen.

### 2.3 Immunocytochemistry

#### Primary cell culture characterisation

Cells were stained for vimentin (fibroblasts, myofibroblasts, endothelial cells; antibody from Progen, Germany), CD31 (endothelial cells; antibody from Pharmingen, USA), and α-smooth muscle actin (αSMA; myofibroblasts, smooth muscle cells; antibody from Abcam, USA). Cells were plated onto sterile glass coverslips, cultured as described above, incubated, and fixed before they reached full confluency. Cell-containing coverslips were washed twice in PBS, incubated in acetone at −20°C for 5 min, and washed again with PBS. Blocking solution containing Polysorbat 20 (“Tween 20”) and foetal calf serum were used to reduce non-specific binding during incubation with primary, and subsequently secondary, antibodies. Nuclei were labelled using Hoechst 33342 nuclear counter stain. Images were acquired with a confocal or a wide-field fluorescence microscope. For quantifying ɑSMA content, a threshold was used to define stained and unstained cells. We used the “Phansalkar” method in ImageJ to create a local threshold based on the minimum and maximum intensities of fluorescence in the proximity of every pixel.^38^ Using this threshold, a macro was designed in ImageJ to identify green fluorescence and count all nuclei marked by Hoechst stain. *Piezo1 and BK_Ca_ immunocytochemistry:* Cells were plated on fibronectin-coated glass coverslips (10 mm diameter), using 24-well plates seeded at 100,000 cells per well. Cells were fixed for 10 min using methanol (5%) and acetic acid at −20°C. Fixed cells were washed with PBS at room temperature (RT) and permeabilized with Triton 0.3% in PBS for 15 min, then incubated during 2 h at RT in the following blocking buffer: PBS containing BSA 4% goat serum 1%, and triton 0.03% for. Primary antibodies: anti-Piezo1 (proteintech, raised in rabbit, 1/300), anti-KCNMA1 (Abnova, Taipei, Taiwan, raised in mouse, 1/500), were incubated 1 hour RT in the blocking buffer. After several washes, secondary antibodies: anti-rabbit IgG AlexaFluor 647 (Invitrogen, Carlsbad, USA, raised in donkey, 1/1250) or anti-mouse IgG AlexaFluor 568 (Invitrogen, raised in donkey, 1/1250) were incubated 50 min at RT. Hoechst 33342 (Molecular Probes, Eugene, USA, 1/5000) were added before mounting with Polyvinyl alcohol mounting medium (with DABCO^®^ antifade reagent, Fluka, Charlotte USA). Images were acquired with LSM 880M (Carl Zeiss, Oberkochen, Germany) with Zen software, maximal z resolution in these conditions was 0.618 μm. Colocalisation was quantified using Pearson’s R coefficient calculated with ImageJ coloc2 plugin.

### 2.4 Electrophysiology

The patch-clamp technique was used to characterise ion channel activity in primary cultures of atrial fibroblasts. Experiments were performed at room temperature (20°C), using a patch-clamp amplifier (200B, Axon Instruments, USA) and a Digidata 1440A interface (Axon Instruments). Recorded currents were digitized at 3 kHz, low-pass filtered at 1 kHz, and analysed with pCLAMP10.3 software (Axon Instruments) and ORIGIN9.1 (OriginLab, USA). Cell-attached patch-clamp recordings were performed using bath and pipette solutions previously described for characterizing Piezo1 channels.^39^ In short: pipette medium contained (in mmol/L): NaCl 150, KCl 5, CaCl_2_ 2, HEPES 10 (pH 7.4 with NaOH); bath medium contained (in mmol/L): KCl 155, EGTA 5, MgCl_2_ 3, and HEPES 10 (pH 7.2 with KOH). Average pipette resistance was 1.3 MΩ. Culture medium was removed and exchanged for the bath solution at least 5 min before the start of electrophysiological measurements to wash-out culture medium and streptomycin (a blocker of SAC).^40^ Membrane patches were stimulated with brief (500 ms) negative pressure pulses of increasing amplitude (from 0 up to −80 mmHg, in 10 mmHg increments unless otherwise stated), applied through the recording electrode using a pressure-clamp device (ALA High Speed Pressure Clamp-1 system; ALA Scientific, USA). To confirm channel identity, we applied the spider toxin peptide *Grammostola spatulata* mechanotoxin 4 (GsMTx4) L-isomer (10 μmol/L, H_2_0 as solvent, CSBio, Menlo Park, CA, USA), a known blocker of cation non-selective SAC, including Piezo1^41^ and potassium-selective ion channels such as BK_Ca_.^42^ The holding voltage for all experiments was −80 mV when recording Piezo1. Current-pressure curves were fitted with a standard Boltzmann function 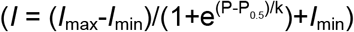, where *I*_max_ is the highest value of current during a pressure pulse, *I*_min_ is the lowest one, and P_0.5_ is the pressure required to obtain half-maximal activation and k is the time constant.

BK_Ca_ channel activity in the absence of additional mechanical perturbation was recorded by depolarising the membrane (from −10 mV up to +60 mV, in 10 mV increments), holding each potential for 22 s. In some cases, the membrane was depolarised up to +80 mV for illustration purposes. The fraction of time during which a channel was open was measured to define the open probability of the channel. To confirm channel identity, the BK_Ca_ activator NS11021 (10 μmol/L or 5 μmol/L, courtesy of Bo Bentzen, Denmark) as well as the BK_Ca_ blockers paxilline (3 μmol/L with 0.03% DMSO, Sigma-Aldrich, Darmstadt, Germany) and iberiotoxin (100 nmol/L, Tocris, Bristol, UK) were used. To assess stretch responses of BK_Ca_, membrane patches were held at +50 mV and stimulated with long (10 s) negative pressure pulses of increasing amplitude (10 mmHg increments). To assess the possible relevance of extracellular calcium on BK_Ca_ activation during stretch, a calcium-free pipette solution was used. It contained (in mmol/L): NaCl 150, KCl 5, EGTA 6, HEPES 10 (pH 7.4 with NaOH). To further confirm channel identity, the dependence of BK_Ca_ channel activity on intracellular Ca^2+^ concentration was measured in the inside-out patch-clamp configuration. Patch excision was achieved by a fast upward displacement of the patch pipette. The bath medium (a nominally Ca^2+^-free solution in these experiments) was supplemented with CaCl_2_ to obtain Ca^2+^ concentrations of 1, 10 and 10 μmol/L.

In order to probe possible interactions between Piezo1 and BK_Ca_ activity, a protocol was used that first activated Piezo1 with a negative pressure pulse (−50 mmHg for 1 s) while holding the patch at −80 mV, and subsequently BK_Ca_ by releasing pressure back to 0 mmHg while clamping the patch to +50 mV (see Fig. 5A for illustration).

Gigaohm seal resistance was systematically checked before and after each protocol: seals having a resistance below 1 GΩ were rejected. If following a stretch protocol, the current did not return to baseline within 20 s, the recording was rejected.

### 2.5 Molecular biology

All reagents, kits and instruments used for molecular biology analysis were supplied by Thermo Fisher Scientific, Germany.

#### mRNA expression levels

Isolation of total RNA was performed using TRIzol Reagent, and frozen samples (made from freshly isolated cells, or from cultured cells for knockdown experiments) were processed according to the manufacturer’s protocol. RNA concentration was quantified by spectrophotometry (ND-1000, Thermo Fisher Scientific, Germany) and synthesis of single-stranded cDNA was carried out as reported before^43^ with the Maxima First Strand cDNA Synthesis Kit, using 3 μg of total RNA. Quantitative real-time polymerase chain reactions (RT-qPCR) was performed as described earlier.^43^ Briefly, 10 μL were used per reaction, consisting of 0.5 μL cDNA, 5 μL TaqMan Fast Universal Master Mix and 6-carboxyfluorescein (FAM)-labelled TaqMan probes and primers. Primers were analysed using the StepOnePlus (Applied Biosystems, Foster City, CA, USA) PCR system. The importin-8 housekeeping gene (IPO8), or glyceraldehyde 3-phosphate dehydrogenase (GAPDH), was used for normalisation. All RT-qPCR reactions were performed as triplicates and control experiments in the absence of cDNA were included. Means of triplicates were used for the 2−ΔCt calculation, where 2−ΔCt corresponds to the ratio of mRNA expression *versus* IPO8. Oligonucleotide sequences are available on request.

#### *PCR-based detection of the* Stress-Axis Regulated Exon *(STREX)*

Total RNA was isolated from freshly isolated fibroblasts and reverse transcribed into cDNA as described earlier. Primers were designed to flank the putative STREX sequence (forward: 5’ – CTGTCATGATGACATCACAGATC – 3’; reverse: 5’ – GTCAATCTGATCATTGCCAGG – 3’, Fig. 4E). PCR were performed to amplify the respective sequence from cDNA of isolated cells from 4 patients. PCR amplification was performed for 40 cycles, using the TaqMan Fast Advanced Master Mix (Cat. No. 4444557, ThermoFisher) according to the manufacturer’s instructions. Subsequently, PCR products were separated by agarose gel electrophoresis and ethidium bromide was visualized by an E-BOX VX2 2.0 MP (Peqlab, Erlangen, Germany) documentation system.

#### Silencing of Piezo1

small interference RNA (siRNA)-mediated knockdown was achieved by transfection using HiPerFect (Qiagen, Venlo, The Netherlands), used according to the manufacturer’s instructions. Pools of 4 siRNA were applied at a final concentration of 8 nmol/L for Piezo1 and Piezo2 (SMARTpool, Dharmacon, Lafayette, USA); concentrations of scrambled controls (Dharmacon) were adjusted accordingly. Knockdown efficiency was assessed by RT-qPCR 48 h after transfection, and electrophysiological experiments were performed 72 h after transfection. The empty vector (siNT) and Piezo1 EGFP constructs (siPiezo1) were transfected in primary human atrial fibroblasts using jetPEI (Polyplus transfection, Illkirch, France) at 0.5 μg of plasmid DNA per 35-mm dish containing ≈25,000 cells.

### 2.6 Co-immunoprecipitation

Cells were grown to confluence in 60 mm dishes, washed twice with ice-cold PBS, lysed with immuno-precipitation (IP)-lysis buffer (from IP Lysis kit Pierce) and supplemented with anti-protease (Roche, Basel, Switzerland). Co-immuno-precipitation (Co-IP) were done following the manufacturer protocol (Pierce co-IP kit). For antibody immobilisation, 10 μg of mouse monoclonal anti-Piezo1 (MyBioSource, San Diego, USA), mouse anti-KCNMA1 (Abnova), mouse monoclonal anti-Na^+^/K^+^-ATPase β1 subunit (Sigma) were added to 50 μL of AminoLink Plus Coupling Resin (Creative Biolabs, New York, USA). Control was done with anti-mouse IgG (Sigma).

Co-immunoprecipitation eluates were subjected to 8% sodium dodecyl sulphate– polyacrylamide gel electrophoresis (SDS-PAGE) before transfer onto a Polyvinylidene fluoride membrane. Membrane was saturated with 5% low-fat milk in Tris-buffered saline containing tween 0.1% during 1 hour at RT and then probed with primary antibodies: anti-Piezo1 (rabbit, Proteintech, 1/1000), anti-KCNMA1 (rabbit, Bethyl, 1/1000), anti-Na^+^/K^+^-ATPase ß1 subunit (Sigma, 1/1000), over-night at 4°C. Horseradish Peroxidase conjuguated anti-mouse IgG (Dako, 1/5000) and anti-rabbit IgG (Dako, 1/2000) were used as secondary antibodies. Blots were revealed with Enhanced Chemoluminescence (Millipore) reaction on a Fusion FX7 Edge 2019. Quantification was made with ImageJ software on 8-bit images after background subtraction (50 pixels rolling ball radius).

### 2.7 Statistical analysis

Unless otherwise indicated, values are expressed as mean ± SEM. *N*-numbers refer to the number of tissue donors, *n*-numbers to the number of cells assessed. Differences between groups with n ≥ 21 were evaluated by Student’s *t*-test. For conditions with n < 21, significance of the difference between means was tested with the nonparametric Mann-Whitney test. The Pearson’s correlation coefficient was used to compare Piezo1 and BK_Ca_ localisation. A p-value < 0.05 was taken to indicate a significant difference between means. Designation of significance: * p < 0.05; ** p < 0.01; *** p < 0.001; ns = not significant.

## 3. Results

### 3.1 Fibroblasts and myofibroblasts are the main constituents of right atrial outgrowth cell cultures

Cells obtained by the outgrowth technique from human right atrial appendage did not contain any cardiomyocytes. The majority (98%) of cells stained positive for vimentin, and 1% of the cells were positive for the endothelial cell marker CD31. Antibody functionality was verified in positive controls using human umbilical vein endothelial cell (HUVEC; Fig. S1A-B). Overall, results confirmed that the outgrowth technique yields predominantly fibroblasts-like cells with negligible contamination by endothelial cells. Vimentin-positive cells formed a mixed population of myofibroblasts and fibroblasts. Myofibroblasts constituted an average of 17.9 ± 9.4% of all cells analysed (n > 37,000 cells / N = 11 SR patients), based on ɑSMA staining (Fig. S1C).

### 3.2 Piezo1 in right atrial fibroblasts of patients in SR

In cell-attached patch clamp recordings (holding potential −80 mV), SAC were observed in response to negative pressure pulses in the patch pipette (Fig. 1A). In this particular cell, inward current activated rather slowly at −40 mmHg, whereas at −80 mmHg, the current peaked rapidly and partially inactivated. Activation and inactivation patterns were variable, therefore, in addition to peak-current amplitude, the average current (mean current calculated over the duration of the pulse of pressure) was analysed. Current-pressure curves had a sigmoidal shape, suggesting saturation at patch pipette pressures more negative than −60 mmHg (Fig. 1B). In n = 110 cells from N = 11 patients in SR, the peak-current amplitude was −37.4 ± 2.8 pA at −60 mmHg and the average current amplitude was −19.9 ± 1.7 pA. The current-voltage (I-V) relationship obtained from single channel activity (n = 9 cells from N = 3 SR patients) was linear, with a slope that yielded a channel conductance of 34.2 ± 1.6 pS (activity recorded at −30 mmHg pipette pressure or during deactivation, Fig. 1C). The reversal potential near 0 mV suggests the presence of a cation-nonselective SAC in human atrial fibroblasts. This was further tested by employing the blocker GsMTx4, which inhibited SAC activity: the average current at −60 mmHg was −42.5 ± 8.3 pA (n = 21; N = 3) in control conditions, *versus* −10.4 ± 2.0 pA (n = 24; N = 3) in the presence of GsMTx4 (Fig. 1D).

**Figure 1:**
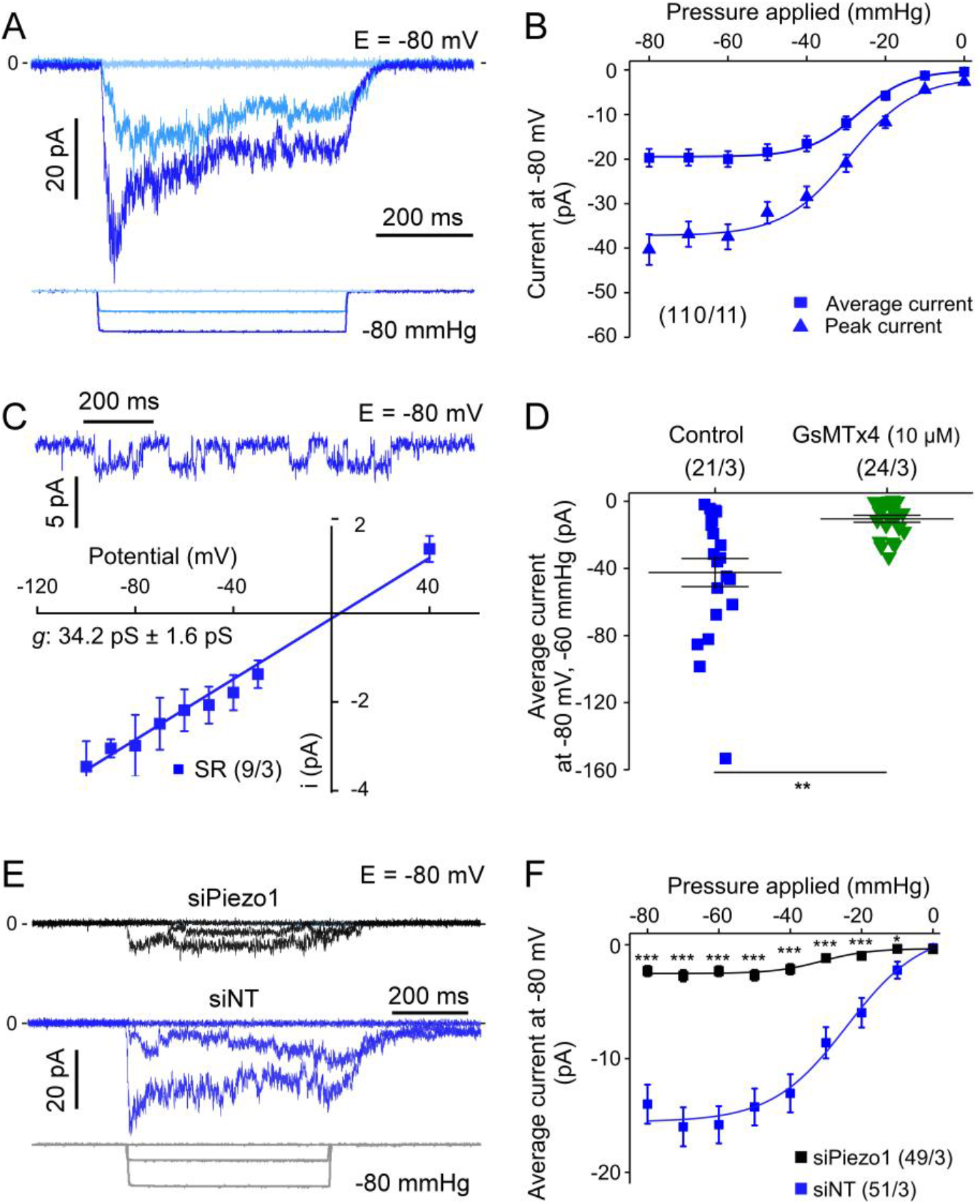
Cation non-selective SAC activity, compatible with Piezo1, is present in human right atrial fibroblasts from patients in sinus rhythm (SR), including cells from passages 0 to 4. **A**: SAC activity, elicited by pulses of negative pressure in cell-attached mode. **B**: Average and peak-currents for all negative pipette pressures tested (from 0 to −80 mmHg); numbers in brackets state n of cells (here 110) and N of tissue donors (here 11) throughout all illustrations. **C**: Top: Single SAC channel activity, activated by a −30 mmHg pressure pulse. Bottom: I-V curve for single channel currents recorded at −30 mmHg or during deactivation, the slope of the straight line was calculated by linear regression (conductance g = 34.2 ± 1.6 pS). **D**: SAC activity at −60 mmHg under control conditions and with GsMTx4 L-isomer (10 μmol/L) in the pipette solution (same patients used for the two conditions). **E**: Representative traces of SAC activity with siRNA targeting Piezo1 (siPiezo1; black trace) and in presence of a non-targeting siRNA (siNT; blue trace), and. **F**: Summary of pressure-effects on average SAC current in cells transfected either with siNT or with a pool of 4 siRNA directed against Piezo1. All recordings, except for I-V curve, were obtained at −80 mV. For all figures: asterisks indicate statistical significance, statistical analysis is described in the section 2.7.

In order to test which SAC contributes to the observed current, Piezo1 was knocked down using a pool of siRNA. Under these conditions, stretch-induced current activity was strongly reduced at all pressure levels tested (Fig. 1E-F); at −60 mmHg, the average current was −15.8 ± 1.6 pA (n = 51; N = 3) in control cells transfected with non-targeting siRNA (Fig. S2A), *versus* −2.3 ± 0.4 pA in siPiezo1 transfected cells (n = 49; N = 3). These values correspond well with the observed 90% reduction in mRNA expression of Piezo1 (but not Piezo2) in siPiezo1-treated cells (Fig. S2B).

Altogether, these results indicate that the SAC activity recorded at −80 mV in human atrial fibroblasts is carried largely by Piezo1.

### 3.3 BK_Ca_ activity in right atrial fibroblasts of patients in SR

In cell-attached voltage clamp experiments without additional suction applied to the membrane, we recorded a distinct ion channel activity at voltages positive to +10 mV. The open probability of this outward current was strongly enhanced by increasing membrane depolarisation (Fig. 2A-B). In addition to this pronounced voltage dependency, the conductance of channels was large, at 125.1 ± 4.9 pS (n = 60; N = 8; Fig. 2C). Based on these biophysical properties, we identified BK_Ca_ as a possible candidate underlying the current.

**Figure 2:**
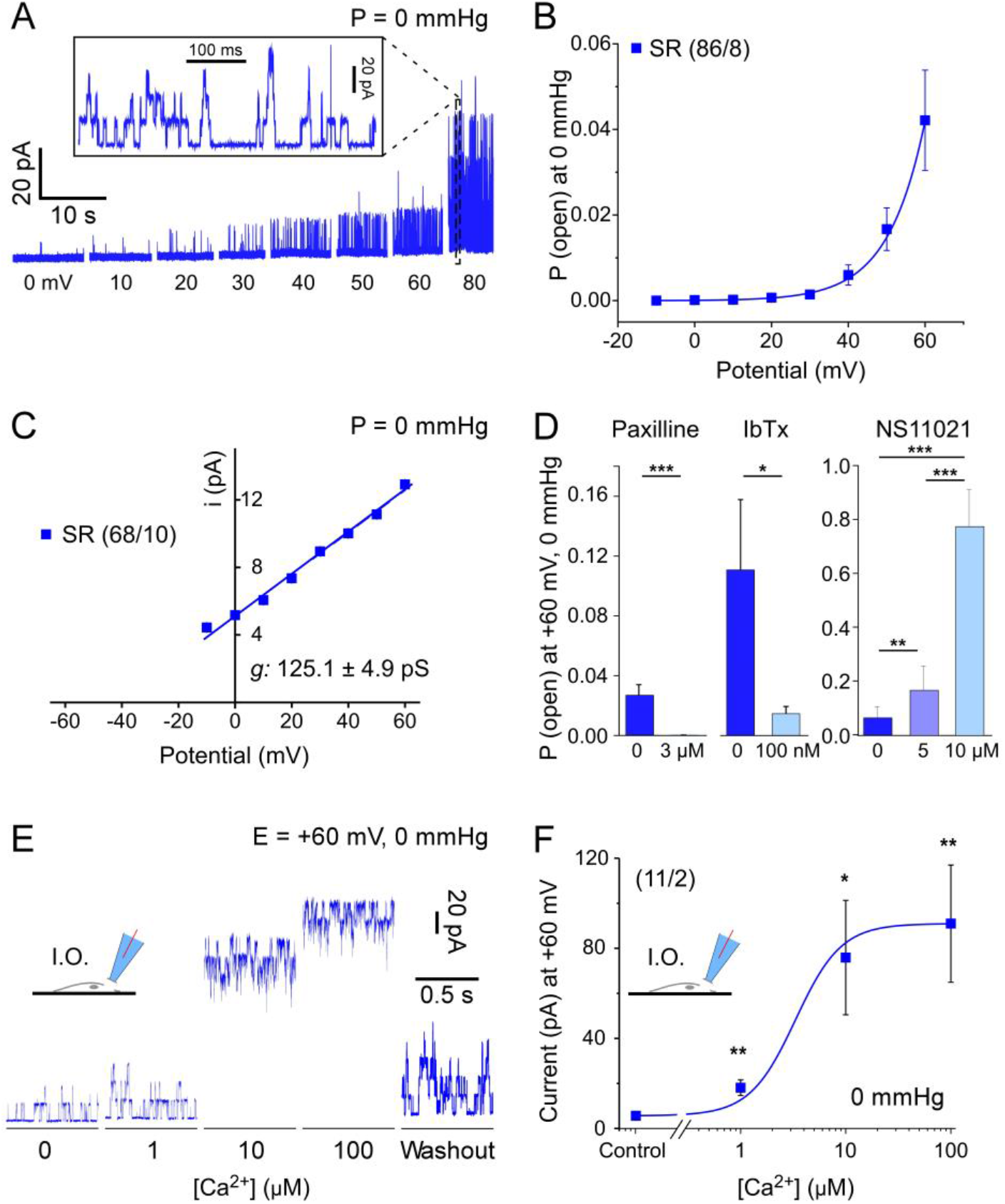
Characterization of BK_Ca_ currents in right atrial fibroblasts from patients in SR. **A**: Current traces in cell-attached patch clamp mode at different holding potentials in the absence of additional mechanical stimulation. Inset: expanded time scale for the trace at +80 mV. **B**: Open probability of BK_Ca_ channels at different potentials from −10 to +60 mV. **C**: I-V curve for single channel currents; straight line slope was calculated by linear regression (conductance g = 125.1 ± 4.9 pS). **D**: Open probability of single channels in control conditions and with 3 μmol/L paxilline (n = 26, N = 2 and n = 21, N = 2 respectively [same SR patients]); in control conditions and with iberiotoxin (IbTx) 100 nmol/L (n = 12, N = 2 and n = 15, N = 2 respectively [same SR patients]); and in control conditions with the BK_Ca_ channel activator NS11021 at 5 and 10 μmol/L (n = 14, N = 2; n = 11, N = 1; n = 18, N = 2, respectively [same SR patients]). Of note, the two controls used for paxilline and iberiotoxin (different patients) illustrate inter-patient variability. **E**: Original recording of BK_Ca_ channel activity at +60 mV in the inside-out configuration with increasing concentrations of Ca^2+^ applied to the cytosolic side of the membrane. **F**: Corresponding quantification of BK_Ca_ channel activity.

As means of further validation, we used a pharmacological approach utilising drugs known to either inhibit^44–46^ or activate^47^ BK_Ca_. Paxilline and iberiotoxin strongly reduced the observed activity (Fig. 2D), while NS11021 caused a robust increase in open probability. The increase in open probability was attributable to an increase in the number of events and dwell time (see Fig S3A and B). There was no significant difference in the conductance of the channel in the presence or absence of NS11021 (Fig. S3C). Interestingly, NS11021 caused a shift in potential dependence of dwell time to less positive potential; this shift was even larger than that in open probability (Fig. S3B).

A key feature of BK_Ca_ channels is their Ca^2+^ sensitivity. We therefore tested the effects of various internal Ca^2+^ concentrations on channel activity (Fig. 2E and F). The inside-out patch configuration was used to expose the cytosolic side of the plasma membrane to increasing Ca^2+^ concentrations, ranging from a nominally Ca^2+^-free environment to 100 μmol/L (pipette was Ca^2+^-free). Upon increase of the Ca^2+^ concentration, a robust activation of the current was observed (from 5.6 ± 1.5 to 90.9 ± 26.0 pA; n = 11, N = 2).

Taken together, our results indicate the presence of BK_Ca_ channel activity in atrial fibroblasts from patients in SR.

### 3.4 Piezo1 and BK_Ca_ activity in right atrial fibroblasts of patients in AF

Piezo1 activity was detected in all AF and SR patients studied. The percentage of cells in which Piezo1 activity was detected was comparable in both patient populations (87%, n = 52, N = 5 in AF; 85%, n = 112, N = 10 in SR). Average Piezo1 current from cells at passage 0 (matched for cell culture time) was significantly higher in right atrial fibroblasts from patients in AF, compared to SR, at negative pressures of −40 mmHg or more (Fig. 3A-B).

**Figure 3:**
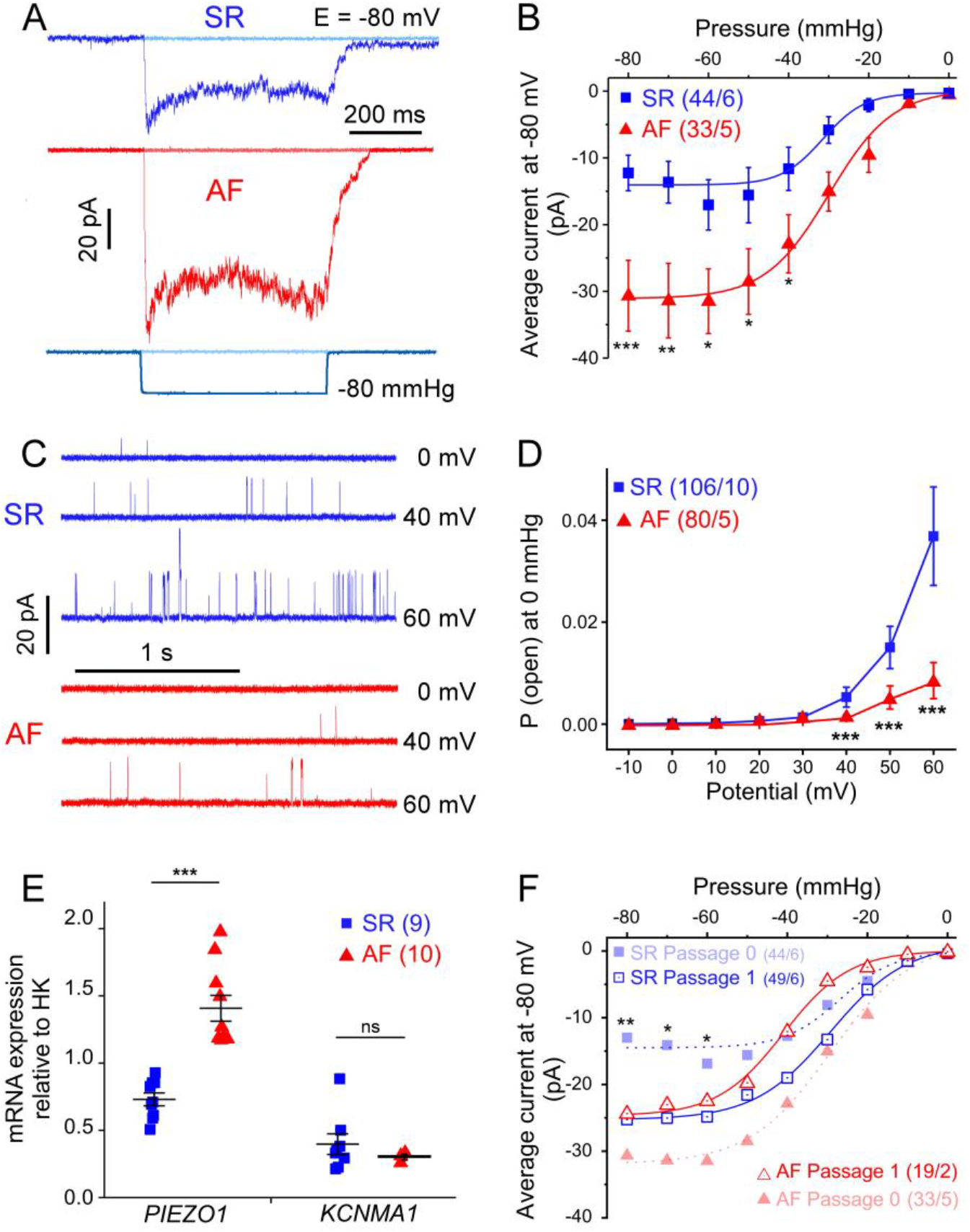
Comparison of Piezo1 and BK_Ca_ channel activity and mRNA expression levels in atrial fibroblasts from patients in SR (blue) and AF (red). **A**: Representative current traces (holding potential −80 mV) activated by 500-ms negative pressure pulse (−80 mmHg). **B**: Mean current-pressure curve for Piezo1, cells from passage 0 only. **C**: Representative traces of single BK_Ca_ channel activity in fibroblasts from an SR and AF patient at different holding potentials. **D**: Voltage dependence of open probability of BK_Ca_ channels from all patches studied. **E**: mRNA expression levels of Piezo1 and KCNMA1, normalised to the housekeeping gene (HK), in freshly isolated non-myocytes from patients in SR and patients in AF. **F**: Current-pressure relationship (average current; holding potential −80 mV) of cells from SR and AF patients at passage 0 (dotted lines; re-plotted for comparison from panel B) and passage 1 (solid lines).

BK_Ca_ activity was also observed in all patients. In some cells, we did not detect BK_Ca_ channel activity under our conditions, and no other channel activity could be measured in these ‘silent’ patches. The fraction of silent patches was larger in cells from AF than SR patients (60% *versus* 34%; n = 80, N = 5 for AF; n = 106, N = 10 for SR). In contrast to Piezo1, BK_Ca_ channels activity was significantly lower in fibroblast from AF than SR patients for voltage clamp steps of +40 mV or more (Fig. 3C-D) even after excluding silent patches (not shown). The reduced open probability was due to a lower number of events in cells from AF tissue; mean dwell times were not significantly different in AF and SR (Fig. S4A-B).

The slopes of the I-V curves, both for Piezo1 or BK_Ca_ channels, were not different in cells from AF and SR patients (Fig. S4C-D). The distribution of cells exhibiting the various activation-inactivation patterns of Piezo1 currents was also not significantly different in cells from AF compared to cells from SR patients (Fig. S4E). These results suggest that the observed differences in Piezo1 and BK_Ca_ activities of fibroblasts from AF tissue, compared to SR, are likely to be be due to altered channel presence, rather than changes in channel properties.

To test the hypothesis that AF is associated with alterations in the expression of Piezo1 and BK_Ca_, we measured the levels of mRNA encoding *PIEZO1* and *KCNMA1*^48, 49^ using RT-qPCR in human right atrial non-myocytes. Freshly isolated non-myocytes, obtained by enzymatic dissociation, were used to capture gene expression levels without any culture time, to be as close as possible to tissue conditions (Fig. 3E). Piezo1 expression levels in non-myocytes from AF patients were almost twice the level of SR cells. For BK_Ca_ channel expression, we did not observe significant differences between the AF and SR groups (Fig. 3E). For control purposes, the purity of the non-myocyte fraction, obtained with our enzymatic dissociation method, was checked by quantifying the expression of typical markers for non-myocytes and cardiomyocytes, *i.e*. vimentin and troponin, respectively (Fig. S5). Expression of vimentin was higher in batch containing isolated non-myocytes than in cardiomyocytes, whereas expression of troponin was higher in isolated myocytes, suggesting that the isolation protocol yielded a significant enrichment of the desired cell type.

Piezo1 activity was found to remodel over culture time (Fig. 3F and S6). Piezo1 activity in cells from SR patients at passage 1 was significantly higher than in passage 0. No further increase of Piezo1 activity was detected at passage 2 (not shown). Cells from AF patients start off with a significantly higher level of Piezo1 than SR cells at passage 0. There is no further significant change in Piezo1 activity in cells from AF tissue (when comparing passage 1 to passage 0), and the initial difference between cells from SR and AF patients is lost at passage 1.

These results indicate that the two channels are differentially regulated in AF, with an increase in Piezo1 activity and expression, and a down-regulation of BK_Ca_ activity.

### 3.5 BK_Ca_ activation during stretch: evidence for two mechanisms

BK_Ca_ channels have been described as mechano-sensitive,^50^ activated directly by membrane stretch.^51^ This has been contrasted by the suggestion that they may also respond to mechanical stimuli indirectly, by sensing changes in intracellular Ca^2+^-concentration, caused by stretch-induced Ca^2+^ and/or Na^+^ entry through other SAC.^52^ Therefore we tested whether BK_Ca_ channel activity in human atrial fibroblasts was modified by stretch.

Fibroblasts from patients in SR were voltage-clamped to +50 mV in cell-attached mode and the patched membrane was simultaneously subjected to negative pressure pulses. The current traces in Fig. 4A show an increase of the typical large-conductance BK_Ca_ channel activity at negative pressures, compatible with stretch-dependent activation. In this particular patch, the maximum number of simultaneously open channels was 3.

**Figure 4:**
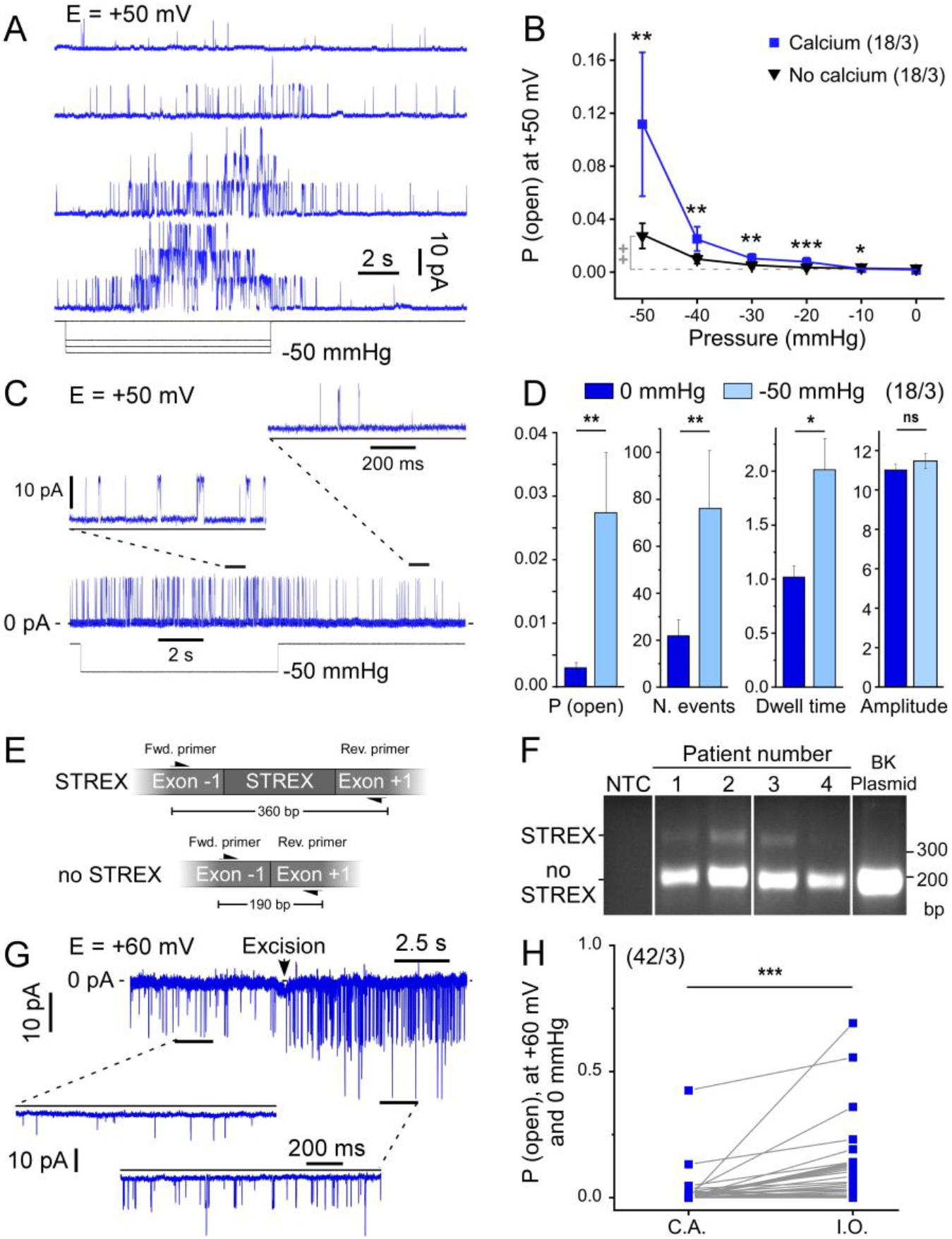
BK_Ca_ stretch-response in human atrial fibroblasts from patients in SR. **A**: Representative current traces at +50 mV in a cell-attached patch from a fibroblast subjected to negative pressure pulses (cell from a donor in SR). **B**: BK_Ca_ open probability in response to stretch with (2 mmol/L) and without Ca^2+^ in the pipette solution. Asterisks indicate statistical significance versus control (with calcium), “+” indicate statistical significance versus no pressure. **C:** Representative recording showing single channel events with (−50 mmHg) and without pressure in a nominally Ca^2+^-free environment pipette solution at +50 mV. **D:** Quantification of the channel open probability, number of single channel events (N. events), dwell time (ms) and single channel amplitude (pA) under conditions described in C. **E:** Schematic representing the position of the two PCR primers used to detect presence or absence of STREX in whole cell mRNA from freshly isolated non-myocytes. bp: base pairs **F:** PCR results showing presence of BK_Ca_ mRNA with and without STREX in 4 patients. NTC: no template control. A purified BK_Ca_ plasmid without STREX is used as a positive control. **G:** Representative recording illustrating the effect of patch excision (from cell-attached to the inside-out configuration) on single channel activity at +60 mV and without pressure; standard Ca^2+^ conditions: nominally Ca^2+^-free bath solution and 2 mmol/L Ca^2+^ in the pipette solution. **H:** Quantification of the channel open probability in the conditions described in G.

The mean open probability of BK_Ca_ channels in the presence of 2 mmol/L Ca^2+^ in the pipette solution was significantly enhanced by increasing negative pressure (shown for n = 18 cells from N = 3 SR patients in Fig. 4B). Interestingly, without Ca^2+^ in the pipette solution, the stretch-dependent increase in BK_Ca_ channel open probability was significantly lower (n = 18 cells from the same N = 3 patients) but not abolished (Fig. B). The stretch-induced increase in open probability resulted mainly from a higher number of single channel events and increased dwell time, while single channel current amplitudes were not significantly different (Fig. 4C and D). As the stress-axis regulated exon (STREX) was described as instrumental for BK_Ca_ stretch-activation, its presence was assessed in freshly isolated human atrial fibroblasts (Fig. 4E and F). PCR results demonstrate that STREX is present in most patients tested, although the BK_Ca_ splice variant without STREX is more abundantly expressed. Upon patch excision, the open probability increased (Fig. 4G and H), suggesting an inhibition of channel activity by the cytoskeleton, as previously shown for BK_Ca._^53^ The data in Fig. 4B suggest that the majority of BK_Ca_-activity during mechanical stimulation is dependent on external Ca^2+^. We hypothesised that Ca^2+^ entry may be secondary to activation of other SAC. We therefore assessed Piezo1 and BK_Ca_ channel crosstalk (Fig. 5). After activating Piezo1 with a 1-s long pulse of negative pressure at a holding potential of −80 mV, suction was terminated and the holding potential switched to +50 mV. This yielded a large outward current that decayed slowly and, once the current amplitude had declined sufficiently, single channel activity could be resolved, compatible with BK_Ca_ activity (see inset in Fig. 5A). Piezo1 activity is known to rundown in response to repeat stimulation,^54^ so when the protocol was repeated, average inward current amplitudes declined dramatically as expected (Fig. 5A-B). Interestingly, the outward current observed upon depolarisation declined in a similar manner. With 0 mmol/L Ca^2+^ in the pipette solution, Piezo1 activity was comparable to control conditions (with calcium), but hardly any BK_Ca_ activity was observed (Fig. 5C-D). In some rare patches in which no SAC current was observed at −50 mmHg, the protocol also failed to yield BK_Ca_ activity (Fig. 5E, BK_Ca_ channel presence was confirmed by activating them, using membrane depolarisation [not shown]). Following the same line of thought, after down-regulation of Piezo1 by siRNA, stretch-dependent BK_Ca_ channel open probability was reduced (Fig. 5F).

**Figure 5:**
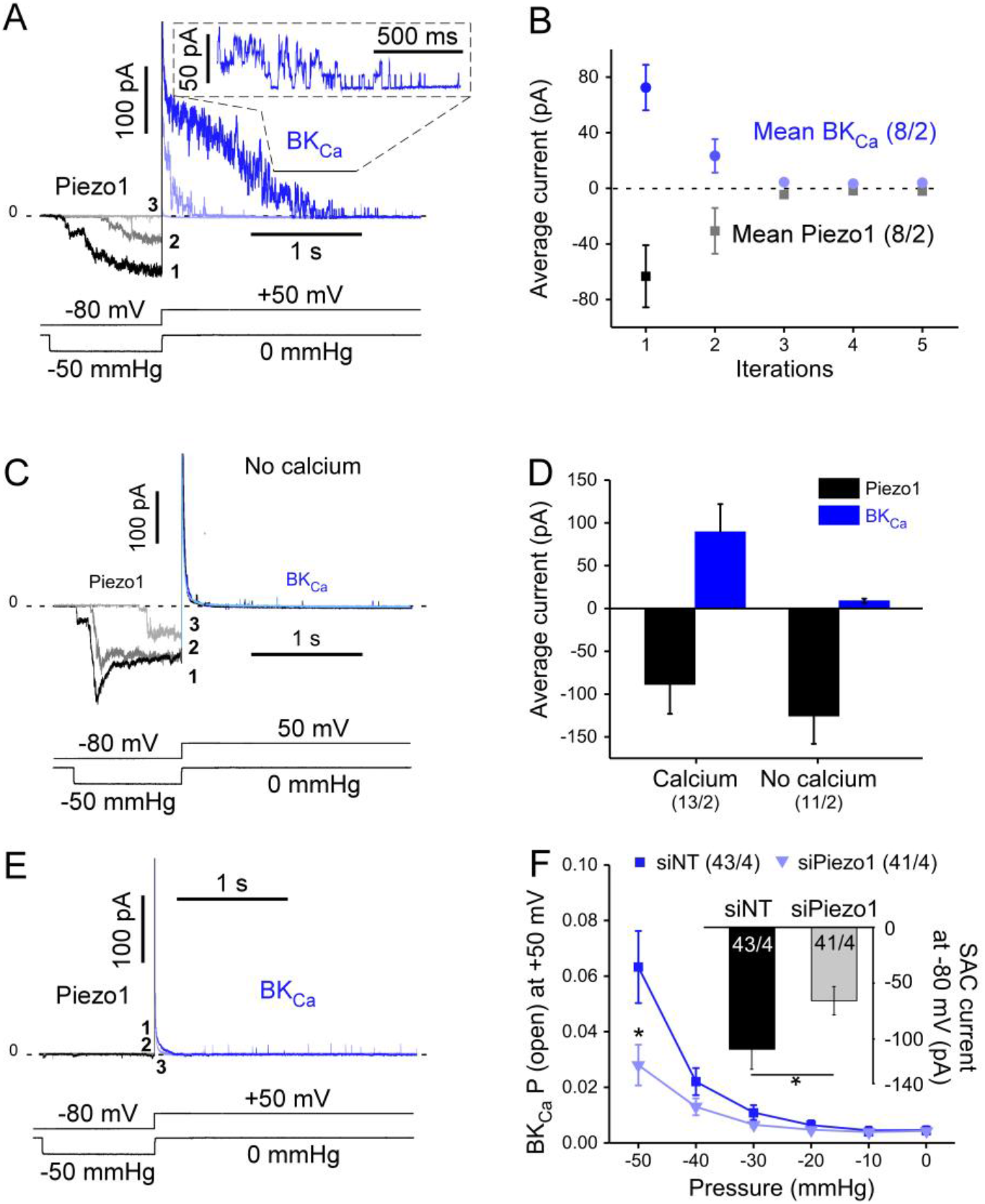
Functional coupling between BK_Ca_ channels and Piezo1 in SR patients. **A**: Piezo1 and BK_Ca_ currents, activated during 3 consecutive sweeps (at 1-min intervals) with the indicated voltage clamp / pressure pulse protocol. The initial suction-induced inward current is carried by Piezo (shades of grey), while the subsequent outward current mainly represents BK_Ca_ (shades of blue). **B**: Mean values of average Piezo1 and BK_Ca_ currents during 5 consecutive runs of the voltage clamp / pressure pulse protocol. **C:** Absence of Ca^2+^ in the pipette solution drastically reduces activation of BK_Ca_ currents during the same voltage clamp / pressure pulse protocol as in A). **D:** Average Piezo1 and BK_Ca_ currents in the presence of 2 mmol/L Ca^2+^ (left) and in a nominally Ca^2+^-free pipette solution (first sweep of the protocol shown in C is quantified. **E:** Lack of BK_Ca_ activation in the rare patches without Piezo1 activity. **F**: Open probability of BK_Ca_ channels (same protocol as in A) in fibroblasts transfected with siRNA targeted against Piezo1 or the empty vector (siNT) or (same patients in SR). Inset: amplitude of average SAC currents, induced by −80 mmHg in the same batch of cells (holding potential −80 mV).

Overall, these results suggest that in human right atrial fibroblasts the apparent mechano-sensitivity of BK_Ca_ channels is mainly secondary to cation non-selective SAC activity, involving Piezo1. The same mechanism was also observed in fibroblasts obtained from AF patients (Fig. S7).

### 3.6 Assessment of Piezo1 - BK_Ca_ structural coupling

To investigate whether the functional interactions of Piezo1 and BK_Ca_ require a physical connection of the two channel proteins, co-immunoprecipitation experiments were performed.

Using either Piezo1 to immunoprecipitate BK_Ca_ or BK_Ca_ to immunoprecipitate Piezo1, no interactions were detected in fibroblast primary cultures (Fig. 6A). Because the quantity of material is limited when working with primary cultures and to improve Piezo1 signal (presence of smear possibly due to post-translational modififications) additional experiments were performed in a human atrial fibroblast cell line.^36^ In these conditions, no interactions between the two channels were observed (Fig. 6B) while the Na^+^/K^+^-ATPase, a known binding-partner of BK_Ca_,^55^ co-immunoprecipitated with BK_Ca_ (Fig. 6C).

**Figure 6:**
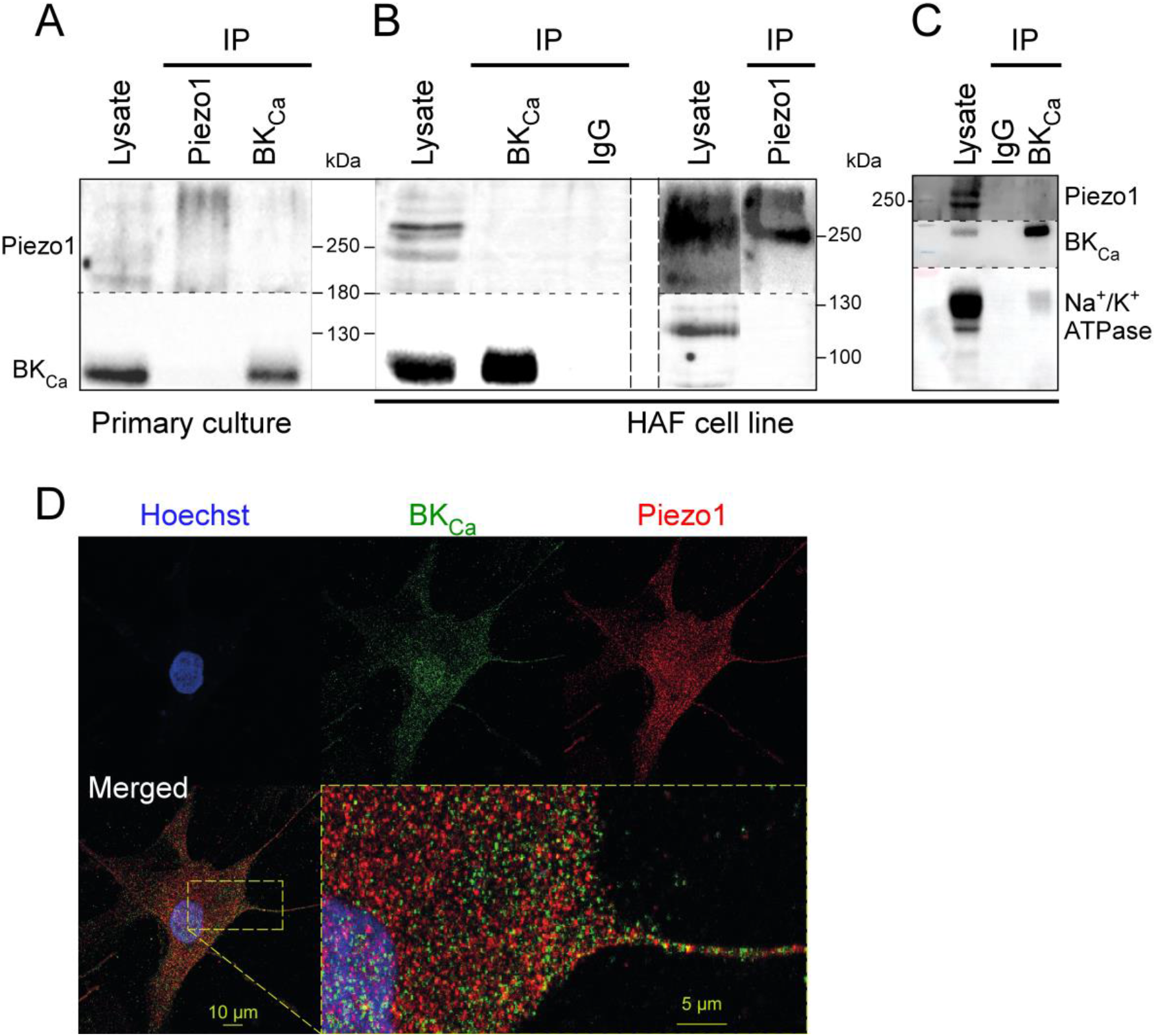
Piezo1 and BK_Ca_ channels do not co-immunoprecipitate and have distinct localisation profiles. **A:** Representative Western blots, showing co-immunoprecipitations with anti-Piezo1 or anti-BK of human atrial fibroblast lysates. **B:** Same experiment performed using a human atrial fibroblast cell line. A column with mouse nonspecific IgG was used as negative control. **C:** Co-immunoprecipitation of Na+/K+-ATPase and BK_Ca_ in the human atrial fibroblast cell line (positive control). **D:** Immuno-staining showing Piezo1 and BK_Ca_ in a primary human right atrial fibroblast.

In agreement with these findings, immunocytochemistry experiments, performed on primary cultures of right atrial fibroblasts from SR patients, revealed distinct localisation profiles of Piezo1 and BK_Ca_ with limited overlap (Fig. 6D). The Pearson’s correlation coefficient of the two signals is 0.46 ± 0.03 (n = 22, N = 1), which indicates no significant degree of correlation. These results suggest that, although Piezo1 and BK_Ca_ are frequently present in the same membrane patches in electrophysiological experiments, they are not physically linked.

## 4. Discussion

We confirm the presence, in human right atrial fibroblasts from patients in SR and AF, of at least two different types of ion currents that are activated during stretch: the cation-nonselective Piezo1 and the potassium-selective BK_Ca_. Our main findings are: (i) activity and expression of Piezo1 are larger in right atrial fibroblasts from AF patients than in cells from SR patients; (ii) activity, but not expression, of BK_Ca_ channels is lower in AF than in SR; and (iii) mechano-sensitivity of BK_Ca_ channels in human atrial fibroblasts is largely secondary to stretch-induced activation of other SAC, including Piezo1.

### 4.1 Cell Identities

As reported in the literature, various cells migrate from small chunks of atrial tissue when placed into appropriate culture medium.^35^ In our hands, this ‘outgrowth technique’ yields 98% vimentin positive cells. In previous work with the same model, we used human fibroblast surface protein as an additional marker to confirm that the vimentin-positive cells are largely fibroblasts.^35^ The endothelial cell marker CD31 was detected in only 1% of our cells, indicating that the majority of the population is of non-endothelial origin. Upon activation, fibroblasts differentiate into myofibroblasts that express αSMA.^56^ The percentage of cells that stained positively for αSMA varied between individuals, with an average of 18% in passage 0 (values ranged from 4% to 38%). In live cell experiments, *i.e*. in the absence of antibody staining, fibroblasts and myofibroblasts could not be differentiated with certainty, based on morphological criteria including size, capacitance and shape. Functional results will therefore reflect a mix of cells, of which ≥80% were fibroblasts.

### 4.2 SAC in human atrial fibroblasts involve Piezo1 channels

In human atrial fibroblasts, voltage-dependent ion channels have been reported,^31, 32, 35^ though relatively little is known about SAC. Negative pressure pulses, applied to the patch pipette at a holding potential of −80 mV, activate inward currents with variable inactivation kinetics. These currents deactivate completely upon pressure release. Using GsMTx4 and a knockdown approach targeting Piezo1, we demonstrate that these SAC in human right atrial fibroblasts are carried largely by Piezo1 (Fig. 1). This result is in line with a recent report by Blythe *et al.*, also on human atrial fibroblasts.^21^

### 4.3 BKCa channels in human atrial fibroblasts

Robust BK_Ca_ channel activity has been reported previously in 88% of human ventricular fibroblasts.^57^ In our study, BK_Ca_ channel activity was present in 40% of right atrial fibroblasts of AF and 66% of SR samples, showing the typical large single channel conductance (125.1 ± 4.9 pS). BK_Ca_ activity was observed in all patients studied. Currents were blocked by paxilline, iberiotoxin and increased in the presence of NS11021. Their open probability was increased both by elevated internal Ca^2+^ concentrations (as previously reported for BK_Ca_)^58^ and by patch excision suggesting sensitivity to cytoskeletal integrity, a known feature of BK_Ca_ channels.^53^ These properties confirm that the observed current is carried by BK_Ca_ channels (Fig. 2).

The study is focussed on right atrial appendage tissue, which is available in the context of open-heart surgery involving extra-corporal circulation, as left atrial tissue is removed more rarely. However, both Piezo1- and BK_Ca_-like activities were confirmed in right and left atrial free wall tissue (Fig. S8). In these cells, single channel amplitudes for Piezo1- and BK_Ca_–like currents were not different from the activity of right auricular fibroblasts included in this paper (not shown), suggesting that the observations reported here are not restricted to right atrial appendage.

### 4.4 Piezo1 and BK_Ca_ currents in human atrial fibroblasts are differentially remodelled during AF

Cells in fibrillating atria are exposed to mechanical loads that may activate SAC. Interestingly, the two channel types investigated here were altered in opposite directions (Fig. 3): whilst presence and stretch-induced activity of Piezo1 were significantly larger in AF compared to SR, consistent with an up-regulation of *PIEZO1* expression, the open probability of BK_Ca_ channels was lower in AF compared to SR, while no differences in expression levels was detected. This change in BK_Ca_ activity in spite of unchanged mRNA levels may be caused by modifications in trafficking, leading to diminished presence of BK_Ca_ channels in the plasma membrane, or an alteration in the regulation of BK_Ca_ gating.

Perhaps surprisingly, cells from SR and AF patients kept their respective phenotype during the first 20-28 days of primary culture. After passaging, however, Piezo1 activity was found to increase in cells from SR patients towards levels that were indistinguishable from AF cells, whilst the intrinsically higher activity in cells from AF patients remained unchanged (Fig. 3F). To avoid effects of prolonged culturing ion channel activity, we focussed our analyses on passage-0 cells. We also attempted to record from freshly isolated fibroblasts. In most cases, repeated negative pressure pulses of significant amplitude (above −20 mmHg) were necessary to obtain seals, in contrast to cultured fibroblasts where seals were generally achieved by a single approach with mild suction (<10 mmHg). As Piezo1 desensitizes with repeat pressure application,^54^ freshly isolated cells were not amenable to obtaining reproducible measurements of SAC activity.

### 4.5 Piezo1 and BK_Ca_ channels are linked functionally, but not structurally

BK_Ca_ channels in human right atrial fibroblasts increased their open probability during mechanical stimulation, applied by stretching the membrane patch in the pipette. Several lines of our experimental evidence suggest that this may, in part at least, be caused by functional crosstalk between stretch-induced activation of Piezo1 and BK_Ca_ activity.

Firstly, we induced transient stretch-activation of Piezo1, followed immediately by recording BK_Ca_ channel activity in the absence of stretch. These experiments identified BK_Ca_ activation as related to the amplitude of the immediately preceding Piezo1 activity (Fig. 5). Desensitisation of Piezo1 during successive activation steps^54^ reduced subsequent BK_Ca_ activity. Similarly, when no Piezo activity was detected in a patch (rare cases), no BK_Ca_ activation was observed either.

Secondly, the functional crosstalk of Piezo1 and BK_Ca_ depends on the presence of Ca^2+^ in the pipette solution: in the absence of extracellular Ca^2+^, Piezo1 currents was still detectable (non-selective cation channel), but BK_Ca_ activation was strongly reduced (Fig. 5C and D). This suggests that a stretch-induced trans-membrane Ca^2+^ flux via Piezo1 may increase in intracellular Ca^2+^ concentration near BK_Ca_ channels and activate them.

Thirdly, knockdown of Piezo1 (see Fig. 5F) reduced the stretch-dependent activation of BK_Ca_. This suppression was incomplete (to about 25% of control). This may be due to partial knockdown of Piezo1 (as shown in the inset of Fig. 5F), or to a contribution from other cation non-selective SAC. A number of other SAC, including Piezo2 (Fig. S2), and canonical transient receptor potential (TRP) channels, including TRPC3 and 6,^59, 60^ are expressed in cardiac fibroblasts, which could influence BK_Ca_ activity. Interestingly, Piezo1 and Piezo2 have similar expression levels (Fig. S2), although the contribution from Piezo2 to the electrical activity recorded after Piezo1 knockdown (Fig. 1F), if any, seems limited.

Taken together, our data suggests that the two channels are functionally coupled. Application of negative pressure to a patch, held at +50 mV, will allow Ca^2+^ entry *via* Piezo1 (the calculated calcium reversal potential, with2 mmol/L Ca^2+^ in the pipette and assuming an intracellular free Ca^2+^ concentration of 100 nmol/L is +125 mV). Similar connections between Ca^2+^-dependent K^+^ channels and Ca^2+^-permeable channels, such as L-type Ca^2+^-channels or SAC, have been described previously,^52, 61, 62^ but so far not in human heart cells. In addition, Piezo1-mediated calcium influx in human atrial fibroblasts has been reported,^33^ which supports our observations.

This functional coupling does not seem to require protein-protein interactions (Fig. 6). It may not depend on the specific pair of proteins studied here – i.e. Piezo1 (and, possibly, other cation non-selective SAC) could influence the activity of other calcium-dependent channels, potentially making them indirectly mechano-sensitive.

Since Piezo1 is non-selective for cations,^22^ Na^+^ will also enter the cell during stretch-activation of Piezo1. Elevated intracellular Na^+^ can increase intracellular Ca^2+^ levels *via* secondary effects, such as mediated by the Na^+^/Ca^2+^ exchanger.^52^ We therefore used Ca^2+^-free conditions, both in the pipette and the bath solution, which would have reduced any contribution by such secondary effects.

While downregulation of Piezo1 and use of Ca^2+^-free conditions strongly reduced BK_Ca_ channel activity, it did not completely abolish it. We interpret the remaining (roughly 25%) BK_Ca_ channel activity as evidence for direct stretch-mediated activation of BK_Ca_ channels. This is supported by the detection of STREX (Fig. 4F) in keeping with previous work.^26, 28^ The lower expression of the STREX-containing variant compared to the BK_Ca_ without STREX corresponds to the electrophysiological observations: the direct stretch-mediated activation represents only 25% of the total BK_Ca_ stretch-induced response (Fig. 4B and F).

### 4.6 Possible functional relevance

The role of, both, the coupling between Piezo1 and BK channels, and the differential remodelling of their activity in the context of AF, for (patho-)physiology of atrial cell and tissue remains to be identified. The resting membrane potentials of fibroblasts isolated from AF and SR patients and cultured on a static substrate did not differ, so functional contributions may need to be explored in the context of cell stretching.

If Piezo1 or/and BK_Ca_ were to alter fibroblast membrane potential during stretch, this could have implications for fibroblasts biology and, possibly, have consequences for electrical excitability, refractoriness, and conduction in myocytes, as fibroblasts and cardiomyocytes can be electrotonically coupled, as demonstrated in murine heart lesions.^63, 64^

We anticipate possible contributions of Piezo1 and BK_Ca_ channels in tissue remodelling as suggested in non AF-related context.^21, 31^ Piezo1 expression and activity have further been proposed to contribute to the control of pro-fibrotic interleukin-6 (IL-6) expression and secretion.^21^ The increased Piezo1 expression and activity reported here in fibroblasts from AF patients, may correspond to an increase in IL-6 signalling. Interestingly, elevated levels of IL-6 correlate with increased left atrial size (as a potential mechanical input),^65^ and AF.^66, 67^

### 4.7 Study limitations and future work

Cells recorded in this study form a mix population of fibroblasts and myofibroblasts. We did not separately quantify the percentage of myofibroblasts *versus* fibroblasts in cells from AF patients. The ratio of myofibroblasts in AF tissue may be higher, compared to control tissue. Further studies will investigate whether fibroblast-myofibroblast phenoconversion influences Piezo1 and BK_Ca_ channels.

Further studies will be required to characterize other essential elements of the BK_Ca_ channel signalling complex, as either or both of the beta or gamma subunits may be changed in the context of AF. This, together with a detailed analysis of BK_Ca_ localisation, would help in revealing why less BK_Ca_ activity is detected in AF fibroblasts while Piezo1 activity is higher.

BK_Ca_ is obviously not the only calcium-activated conductance in fibroblasts. Analysing the effects of Piezo1 opening, for example on the calcium-activated chloride channel anoctamin-1, would be quite relevant in the context of AF, as up-regulation of that channel has been reported to prevent fibrosis after myocardial infarction.^68^

In conclusion, we describe two ion channel populations in human right atrial fibroblasts whose activity is increased during stretch, either directly (Piezo1 and a subset of BK_Ca_ channels) or indirectly (most BK_Ca_ channels, whose activation depends on functional crosstalk with Piezo1). The two channels are differentially regulated in AF, with an increase in Piezo1 activity and expression, and a down-regulation of BK_Ca_ activity. The Patho-physiological relevance of these changes remains to be explored.

## Author Contributions

DJ, AK, SNH, PK, UR and RP contributed to conception, design and interpretation of the study. DJ, AK and ED performed and analysed electrophysiological experiments. DA, EARZ, SP and CS performed and analysed quantitative RT-PCR experiments. DJ and TG performed and analysed immunocytochemical experiments. DA and ASC isolated cells. BA and HG performed and analysed the co-immunoprecipitation and localisation experiments. RE performed the PCR to assess the presence of STREX. SRK provided HAF cells. FB, CS, MK and FAK provided access to surgical tissue samples. DJ, AK, UR and RP drafted the manuscript. All authors contributed to manuscript revision, read and approved the submitted version.

## Acknowledgments

The authors thank all colleagues at the Department for Cardiovascular Surgery of the University Heart Centre Freiburg - Bad Krozingen, and at the CardioVascular BioBank Freiburg, for providing access to human atrial tissue. Special thanks for technical support go to Cinthia Buchmann, Anne Hetkamp, Kristina Kollmar and Gabriele Lechner. We would like to also thank Simone Nübling and Hannah Fürniss for their help concerning patient demographics. We thank Dr Bo Bentzen (University of Copenhagen) for providing us with the compound NS11021. We acknowledge support from SCI-MED for image acquisition and analysis.

