## supplementary figures for "Piezo1 and BK_Ca_ channels in human atrial fibroblasts: interplay and remodelling in atrial fibrillation"

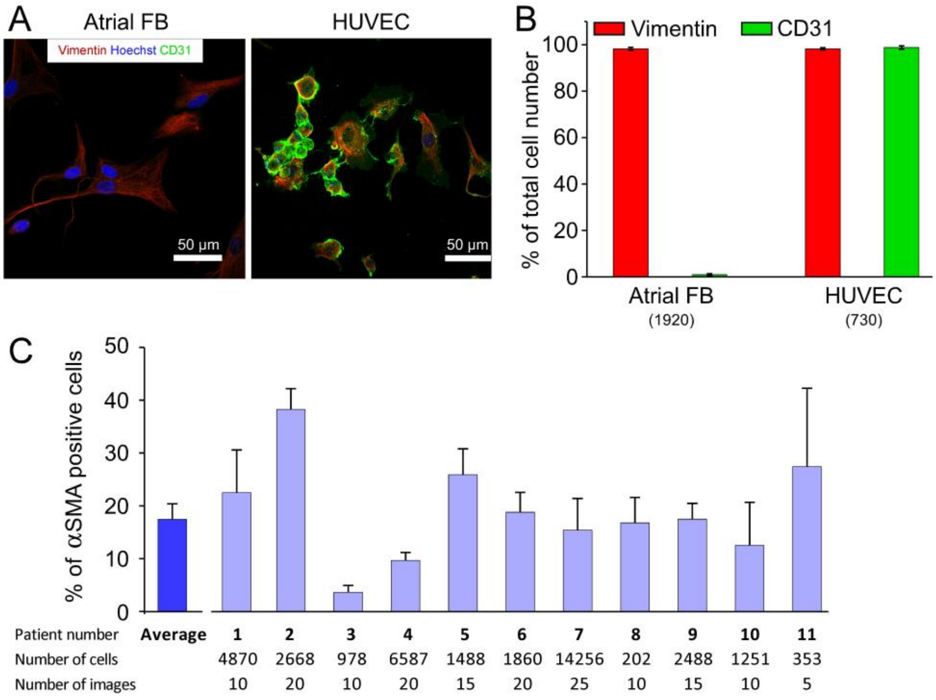


***Supplementary Figure S1**: Composition of non-myocyte population obtained by the outgrowth method. **A**: Example of atrial outgrowth culture and of human umbilical vein endothelial cells (HUVEC, used as a positive control to confirm antibody labelling protocol) stained for vimentin (red), the specific endothelial cell marker CD31 (green), and fibroblast nuclei (blue). **B**: Percentage of vimentin-positive and CD31-positive cells obtained with the outgrowth technique, compared to HUVEC. **C**: Percentage of αSMA expressing cells from N = 11 SR patients (passage 0); mean 17.9 ± 9.4% of n > 37,000 cells analysed.*

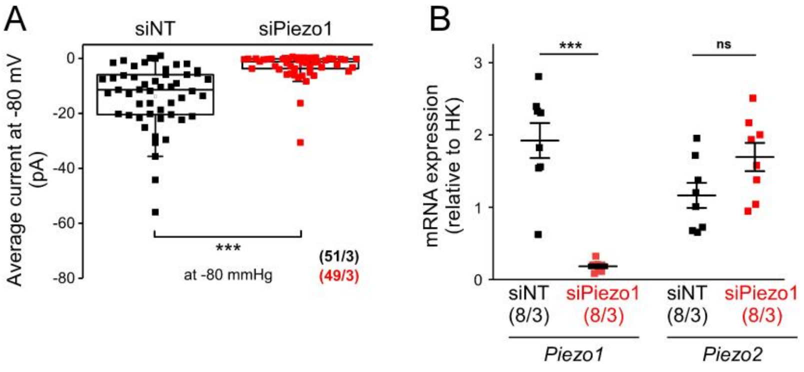


***Supplementary Figure S2**: SAC current as well as mRNA expression (of Piezo1 and Piezo2) in human atrial fibroblasts from N = 3 patients in SR, transfected with siNT or siPiezo1. **A**: Average current at −80 mmHg is significantly reduced after Piezo1 knockdown (holding potential −80 mV). **B**: mRNA expression (relative to housekeeping [HK] gene ß actin.) of Piezo1 and Piezo2 in fibroblasts after transfection with siNT and siPiezo1.*

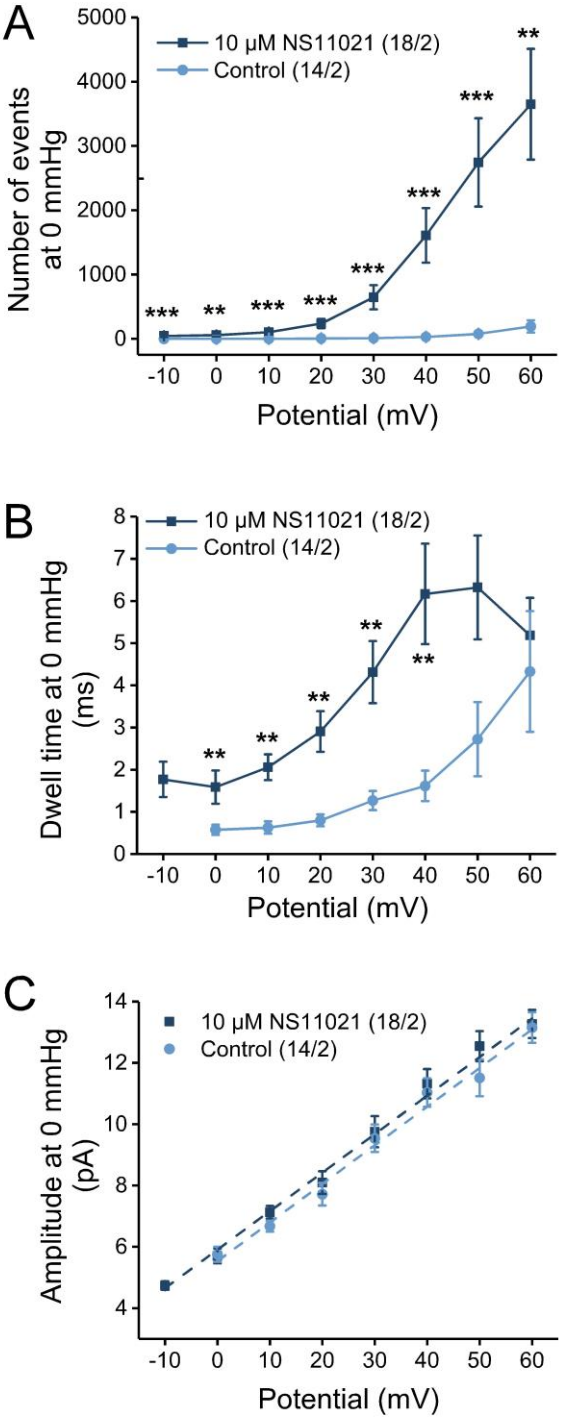


***Supplementary Figure S3: Further characterization of $BK_{Ca}$ channel activity with and without NS11021 (10 µmol/L) in cell-attached membrane patches of human atrial fibroblasts from patients in SR. A****: Number of channel events with and without NS11021 as a function of holding potential.* ***B****: As in A, for dwell time of the channel events with and without NS11021.* ***C****: I-V curves for single $BK_{Ca}$ channel currents in the absence and presence of NS11021 overlap. Statistical analysis is described in the section 2.7*.

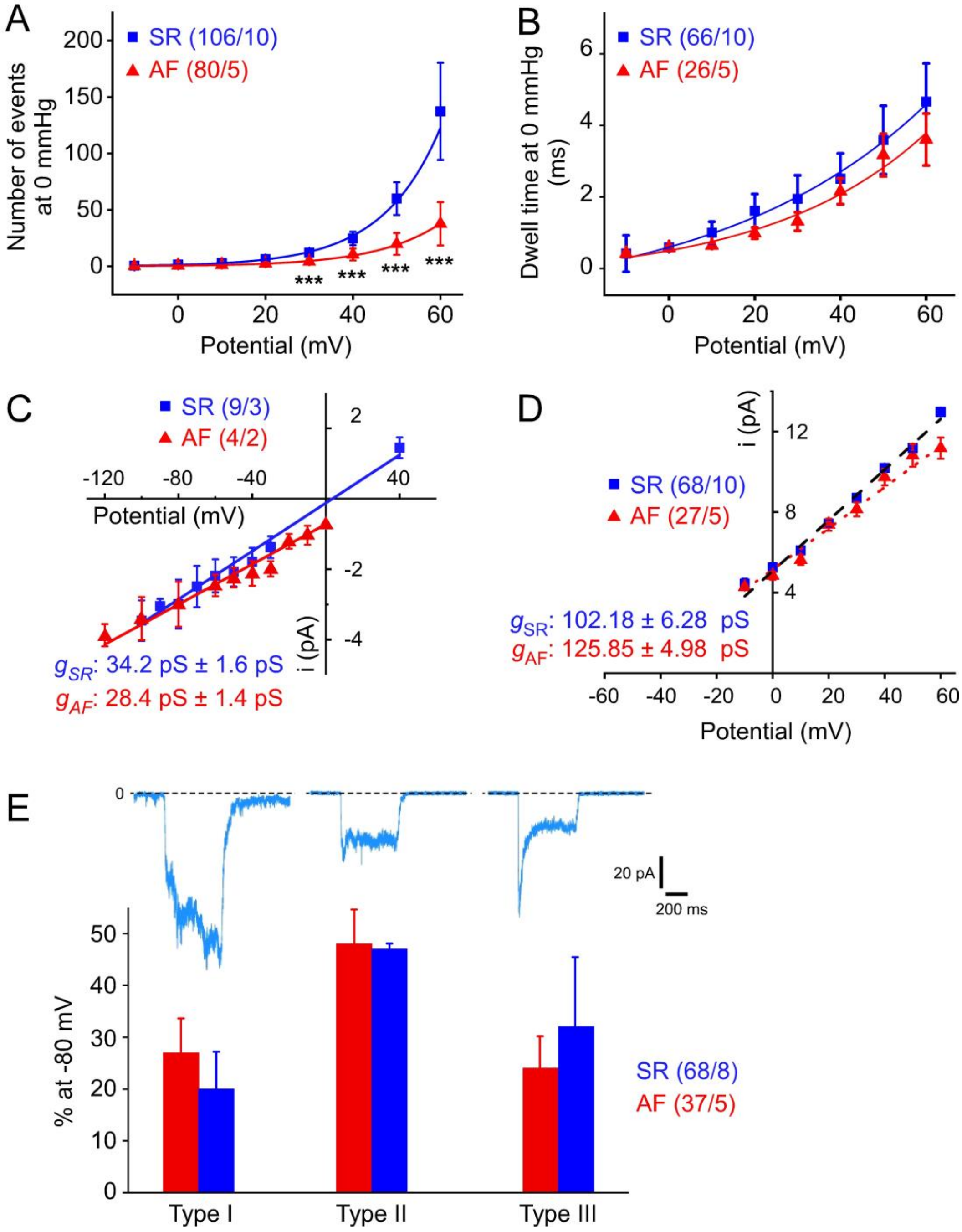


***Supplementary Figure S4: Characteristics of $BK_{Ca}$ and Piezo1 activity in cultured human atrial fibroblasts from patients in SR (blue) and AF (red). A****: Number of $BK_{Ca}$ channel openings as a function of holding potential; **B**: As in A, for dwell time. **C** and **D**: I-V curves for single channel currents of Piezo1 (C) and $BK_{Ca}$ channels (D) in cells from SR and AF. **E**: Examples of 3 distinct activation/inactivation patterns of Piezo1 currents (top), and their relative distribution (bottom) in fibroblasts from AF and SR patients. Currents typically activated very rapidly in response to membrane stretch, followed either (i) by a slower increase in activation and no inactivation (type I); (ii) small inactivation to a steady-state (type II); or (iii) profound inactivation (type III).*

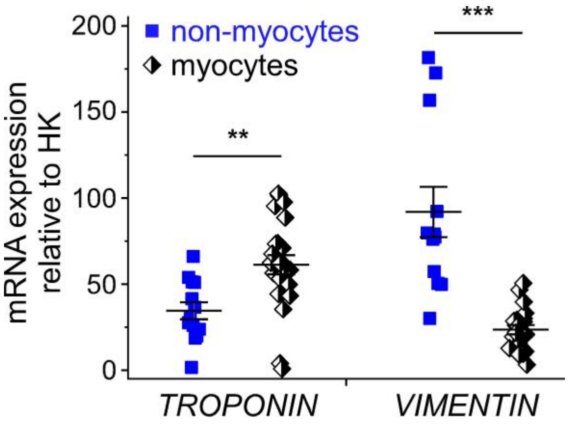


***Supplementary Figure S5: Assessment of cardiomyocyte versus non-myocyte enrichment in cell suspensions used to study mRNA expression.*** *mRNA expression levels of TROPONIN and VIMENTIN, normalised to the housekeeping gene (HK) IPO8, in freshly isolated cells from N = 9 patients in SR, and N = 10 patients in AF. Although cells from 19 patients were used, in some case (especially for the non-mycocyte fraction) the number of data points is less and this is due to an insufficient quantity of material in some samples. Statistical analysis is described in the section 2.7.*

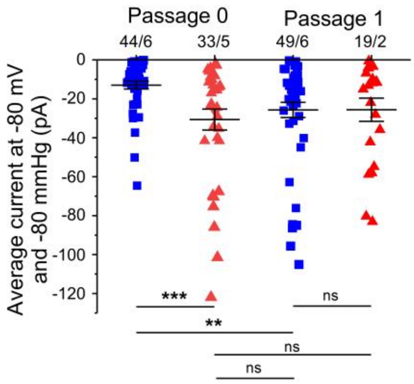


***Supplementary Figure S6: Piezo1 activity in human atrial fibroblasts from patients in SR and in AF at different passages.*** *Average current at −80 mmHg pressure in cells from passage 0 and passage 1 for SR (blue) and AF cells (red). Individual cells are shown. Statistical analysis is described in the section 2.7.*

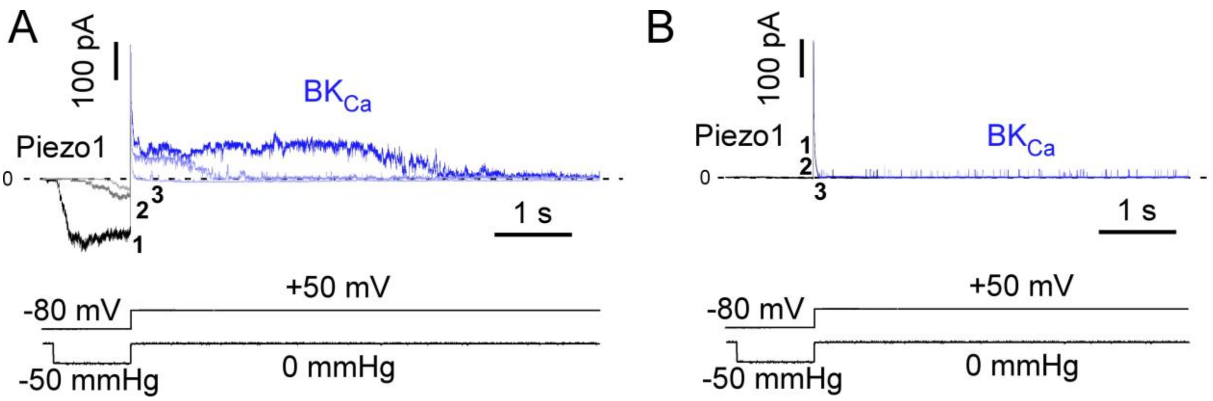


***Supplementary Figure S7: Piezo1 - $BK_{Ca}$ coupling in fibroblasts from AF patients. A****: Representative Piezo1 and $BK_{Ca}$ currents, activated during 3 consecutive sweeps with the indicated voltage clamp / pressure pulse protocol. The initial inward current is carried by Piezo (shades of grey), while the later outward current mainly represents $BK_{Ca}$ (shades of blue).* ***B****: Representative recording illustrating that in the absence of Piezo1 activity pressure does not trigger $BK_{Ca}$ activity although $BK_{Ca}$ channels are present as single channels are observed at +50 mV.*

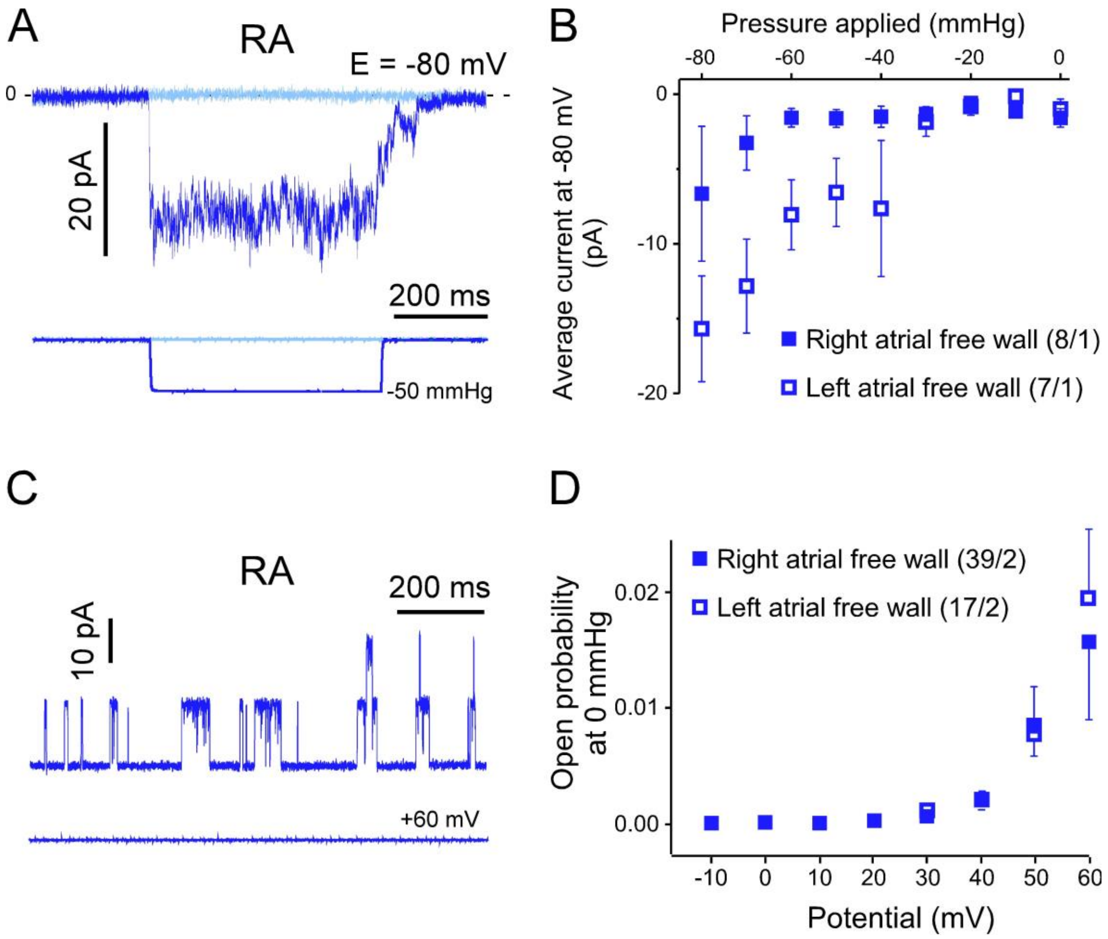


***Supplementary Figure S8: Piezo1- and $BK_{Ca}$-like activities are present in fibroblasts isolated from right and left atrial free wall. A**: Piezo1-like activity, elicited by pulses of negative pressure in cell-attached mode in fibroblasts isolated from the right atrial free wall (RA: right atrium, of which the auriculum used elsewhere is a part). **B**: Average Piezo1-like current for all pressures tested (from 0 to −80 mmHg) for both right and left atrial free wall (n = 8 and 7 cells, respectively; N = 1 per condition). **C**: Top: Large $BK_{Ca}$-like single channel events in fibroblast from right atrial free wall. Bottom: holding potential (+60 mV). **D**: Voltage dependence of open probability of $BK_{Ca}$-like channels in fibroblasts from right and left atrial free wall (n = 39 and 17 cells, respectively; N = 2 per condition).*
